# Granulin loss and TMEM106B risk converge on lysosomal C-terminal fragment pathology in frontotemporal dementia

**DOI:** 10.64898/2026.03.25.713523

**Authors:** Yi Zeng, Jian Xiong, Anastasiia Lovchykova, Thao Phuong Nguyen, Abigail Song, Semma W. Gitler, Monther Abu-Remaileh, Aaron D. Gitler

## Abstract

Frontotemporal dementia (FTD) is the second most common cause of dementia after Alzheimer disease. Mutations in *GRN*, which encodes progranulin, are a major cause of FTD. Common genetic variants in the *TMEM106B* gene modify risk of FTD and the effect is especially strong in *GRN* mutation carriers. Intriguingly, in *GRN* mutation carriers, being homozygous for the protective *TMEM106B* haplotype seems to confer near lifetime protection against FTD. Despite the strong genetic link between *GRN* and *TMEM106B*, how these two genes interact mechanistically has remained unresolved. Recent studies have revealed that a C-terminal fragment of TMEM106B forms amyloid fibrils and accumulates in the brains of older individuals and patients with neurodegenerative disorders, including FTD. How the production of this fragment connects to granulin deficiency is also unknown. Using lysosome immunoprecipitation, we show that granulin deficiency drives the accumulation of the TMEM106B C-terminal fragment within lysosomes in *Grn*-knockout mice and *GRN*-null human iPSC-derived neurons. Recombinant progranulin supplementation reduced TMEM106B C-terminal fragment accumulation. Isogenic neurons carrying the *TMEM106B* risk allele displayed allele-dose-dependent fragment accumulation that was reversible by progranulin. Structural and genetic analyses demonstrated that TMEM106B dimerization stabilizes the protein and limits C-terminal fragment formation. These findings define a lysosomal pathway linking granulin deficiency to TMEM106B C-terminal fragment accumulation and explain how protective *TMEM106B* alleles can confer resistance to FTD, even for *GRN* mutation carriers.

**One Sentence Summary:** Granulin deficiency drives lysosomal accumulation of an amyloidogenic TMEM106B C-terminal fragment, revealing a molecular mechanism that explains how *TMEM106B* alleles can confer risk or protection from frontotemporal dementia.

## Introduction

Frontotemporal dementia (FTD) is a leading cause of early-onset dementia, marked by degeneration of frontal and temporal cortices and pathological inclusions containing either TDP-43 or tau^1–5^. Heterozygous *GRN* mutations cause autosomal-dominant FTD owing to haploinsufficiency of the lysosomal glycoprotein progranulin^2,6^. Complete loss of progranulin causes the lysosomal storage disorder neuronal ceroid lipofuscinosis^7^. Progranulin is proteolytically processed in the lysosome to generate mature peptides called granulins, which play diverse functions in the lysosome^8–10^. How granulin deficiency impairs lysosome function and how this causes neurodegeneration remains unclear.

A genome-wide association study identified polymorphisms in *TMEM106B* as a powerful modifier of FTD disease penetrance^11,12^, particularly among *GRN* mutation carriers^13^. Remarkably, inheriting two copies of the protective *TMEM106B* haplotype nearly abolishes risk of developing disease in some *GRN* mutation carriers^14^. Both gene products localize to lysosomes, yet the mechanistic basis of this striking genetic effect has remained elusive. Initially, it was proposed that genetic variants cause increased risk for FTD by increasing expression of TMEM106B^15,16^. Indeed, *TMEM106B* overexpression perturbs the endolysosomal pathway^17,18^. But, contrary to expectation, *Tmem106b* knockout in mouse exacerbates the *Grn* knockout phenotype^19–21^ and even leads to TDP-43 pathology (the pathological hallmark of FTD caused by *GRN* mutations)^2^. Thus, it remains unclear how *GRN* and *TMEM106B* functionally interact to contribute to FTD risk.

TMEM106B is a type II lysosomal membrane protein involved in trafficking and degradative function^22–24^. Recent cryo-EM studies revealed that TMEM106B can be cleaved to generate a luminal C-terminal fragment that forms amyloid fibrils in aged and diseased brains, including in FTD^24–27^. TMEM106B fibrils predominantly accumulate in neurons and astrocytes and do not appear to colocalize with other amyloid fibrils such as TDP-43, α-synuclein, amyloid-beta, and tau^28–30^. Recent evidence suggests that this cleavage product is not only pathological but reflects physiological turnover–its cleavage and processing are mediated by a series of lysosomal cysteine-type cathepsin proteases^31^. However, how TMEM106B processing in the lysosome connects to *GRN* and *TMEM106B* genetic variants (that confer risk or protection) is unknown.

Using lysosome immunoprecipitation, human iPSC-derived neuronal models, structural analysis, and quantitative peptide analysis in human brain samples, we uncovered a direct molecular link between *GRN* and *TMEM106B*. Our findings reveal how granulin regulates TMEM106B proteostasis, explain the protective effect of *TMEM106B* alleles, and identify dimer stabilization as a potential therapeutic strategy.

## Results

Because of the strong association between genetic variants in *TMEM106B* and risk of FTD, especially in *GRN* mutation carriers^13^, the remarkable ability of some *TMEM106B* variants to confer near lifetime protection against FTD^14^, and the accumulation of TMEM106B C-terminal fragments in FTD patient brain in a risk haplotype-dependent manner^32^, we hypothesized that granulin deficiency might alter lysosomal accumulation of the TMEM106B C-terminal fragment.

To test this hypothesis, we performed lysosome immunoprecipitation (LysoIP)^33^ on granulin-deficient (*GRN* KO) and isogenic wild type (WT) human neurons differentiated from induced pluripotent stem cells (iPSCs). We engineered these cells to stably express LysoTag (lysosomal transmembrane protein 192 (TMEM192) fused to three copies of the human influenza virus hemagglutinin (HA) tag) and confirmed successful purification of intact lysosomes (Figure S1A-C). We used an antibody that specifically recognizes the TMEM106B C-terminal fragment^32^ and found a marked increase in its abundance within lysosomes from *GRN* KO neurons compared to the wild type controls (Figure 1A, B). TMEM106B monomer levels also increased (Figure 1A, B), but the increase in the C-terminal fragment in *GRN* KO neurons was not owing to increased numbers of lysosomes because normalizing to lysosomal-associated membrane protein 1 (LAMP1) or Cathepsin B, a lysosomal lumen protein, still showed increased TMEM106B C-terminal fragment abundance (Figure 1A-B, Figure S1C). We also confirmed the increase of lysosomal C-terminal fragments in *GRN* KO neurons using a different antibody specific to the cleaved C-terminal fragment (Figure S1C). *TMEM106B* mRNA levels did not differ between *GRN* WT and *GRN* KO neurons (Figure 1C), indicating that the impact of granulin deficiency on TMEM106B was at the protein level. We detected increased levels of TMEM106B C-terminal fragment in total cell lysates (Figure 1B, bottom panel), but the effect was much more apparent in the LysoIP samples (Figure 1B, top panel), indicating that granulin deficiency results in lysosomal TMEM106B C-terminal fragment accumulation.

**Figure 1.**
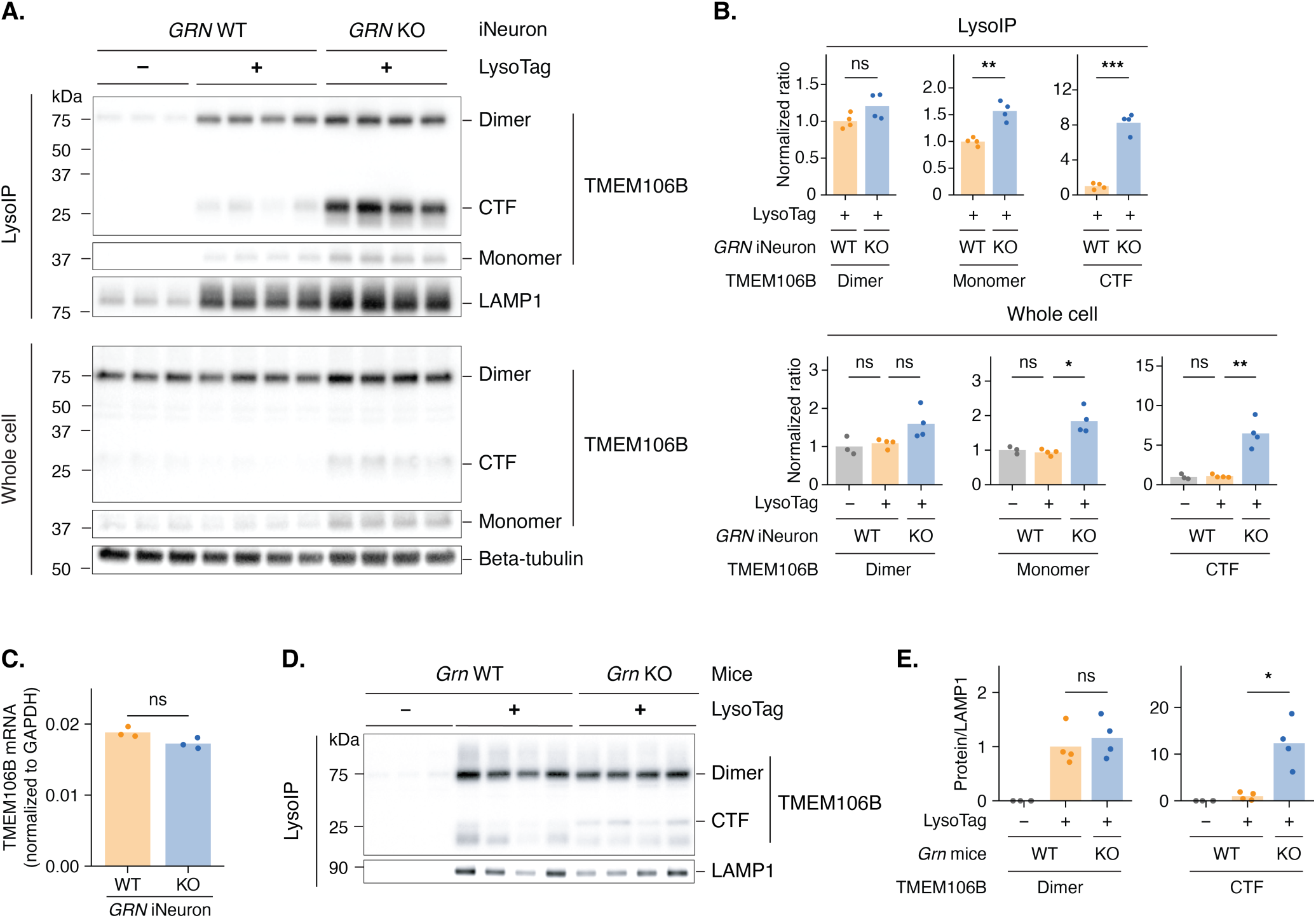
Granulin deficiency leads to the accumulation of TMEM106B C-terminal fragments in the lysosome. A. Western blots show that *GRN* knockout (KO) in iNeurons leads to accumulation of TMEM106B C-terminal fragments (CTFs) in lysosomes. Purified lysosomes (LysoIP; top) and whole cell lysates (bottom) were from *GRN* wild type (WT) iNeurons without LysoTag (C-terminally tagged TMEM192-3xHA), *GRN* WT iNeurons with LysoTag, and *GRN* KO iNeurons with LysoTag. Lysosomes were purified by immunoprecipitation using the LysoTag. B. Quantification of TMEM106B dimers, monomers, and CTFs from panel A. Normalized ratios were calculated by dividing the intensity of each TMEM106B species (dimer, monomer, or CTF) by the loading control (Beta-tubulin for whole cell lysates; LAMP1 for purified lysosomes), then normalizing to the first bar (WT with LysoTag for purified lysosomes or WT without LysoTag for whole cell lysates). C. qRT-PCR analysis shows that *GRN* KO does not alter TMEM106B mRNA levels in iNeurons. D. Western blots show that *Grn* KO in mice leads to accumulation of Tmem106b CTFs in lysosomes. Purified lysosomes were from livers of 6-month-old *Grn* WT mice without LysoTag, *Grn* WT mice with LysoTag, and *Grn* KO mice with LysoTag. Lysosomes were purified by immunoprecipitation using the LysoTag. E. Quantification of Tmem106b dimers and CTFs from panel D. Normalized ratios were calculated by dividing the intensity of each Tmem106b species (dimer or CTF) by the loading control (LAMP1), then normalizing to that of Grn WT with LysoTag. Bar plots represent the mean, and each dot represents a replicate (n = 3-4 replicates per condition). Statistical significance was determined by a two-sided Welch’s t-test: ns (not significant), p > 0.05; *, p ≤ 0.05; **, p ≤ 0.01; ***, p ≤ 0.001; ****, p ≤ 0.0001.

To extend this finding, we examined whether granulin deficiency leads to increased TMEM106B C-terminal fragments *in vivo*. We crossed a transgenic mouse line that constitutively expresses the LysoTag in all tissues^34^ to *Grn* knockout mice^35^. Lysosomes derived from *Grn* knockout mice revealed that granulin deficiency resulted in an increase in endogenous lysosomal Tmem106b C-terminal fragment accumulation compared to those of the WT controls (Figure 1D, E). Thus, granulin deficiency leads to TMEM106B misprocessing and C-terminal fragment accumulation in the lysosome *in vivo*.

Because granulin deficiency increased C-terminal fragment levels without significantly altering dimer levels (Figure 1), we hypothesized that granulin acts downstream of TMEM106B dimer formation to directly regulate C-terminal fragment levels. To test this hypothesis, we first needed a way to isolate the effect of granulin on C-terminal fragments independent of TMEM106B dimerization. Previous evidence suggested that monomers are more prone to cleavage than dimers: in human FTD brain, the *TMEM106B* risk haplotype associates with less dimer and more TMEM106B C-terminal fragments, whereas the protective haplotype associates with more dimer and less C-terminal fragments^32^. Therefore, we engineered TMEM106B mutants that cannot dimerize. Using AlphaFold3^36^ to model the TMEM106B homodimer structure, we identified a conserved zinc-coordination interface where two pairs of cysteine residues from CXXC motifs (one from each monomer) appear to coordinate an intermolecular zinc ion to stabilize the dimer (Figure 2A), similar to the role of the Rad50 zinc hook motif^37^ and a recent finding^38^. We engineered specific mutations to disrupt this interface (CPTC61-64→AAAA; Figure 2B), with or without disrupting a predicted intermolecular disulfide bond (C105A). Both mutants (Allmut and Nmut) abolished dimerization (Figure 2C) and increased C-terminal fragment accumulation (Figure 2C), an effect independent of the N-terminal tag used (Figure 2C, D). Purified lysosomes from Allmut expressing HEK293 cells further confirmed that TMEM106B C-terminal fragments accumulated in lysosomes (Figure 2E, F). These results provide evidence that dimer formation protects TMEM106B from aberrant proteolytic cleavage within lysosomes.

**Figure 2.**
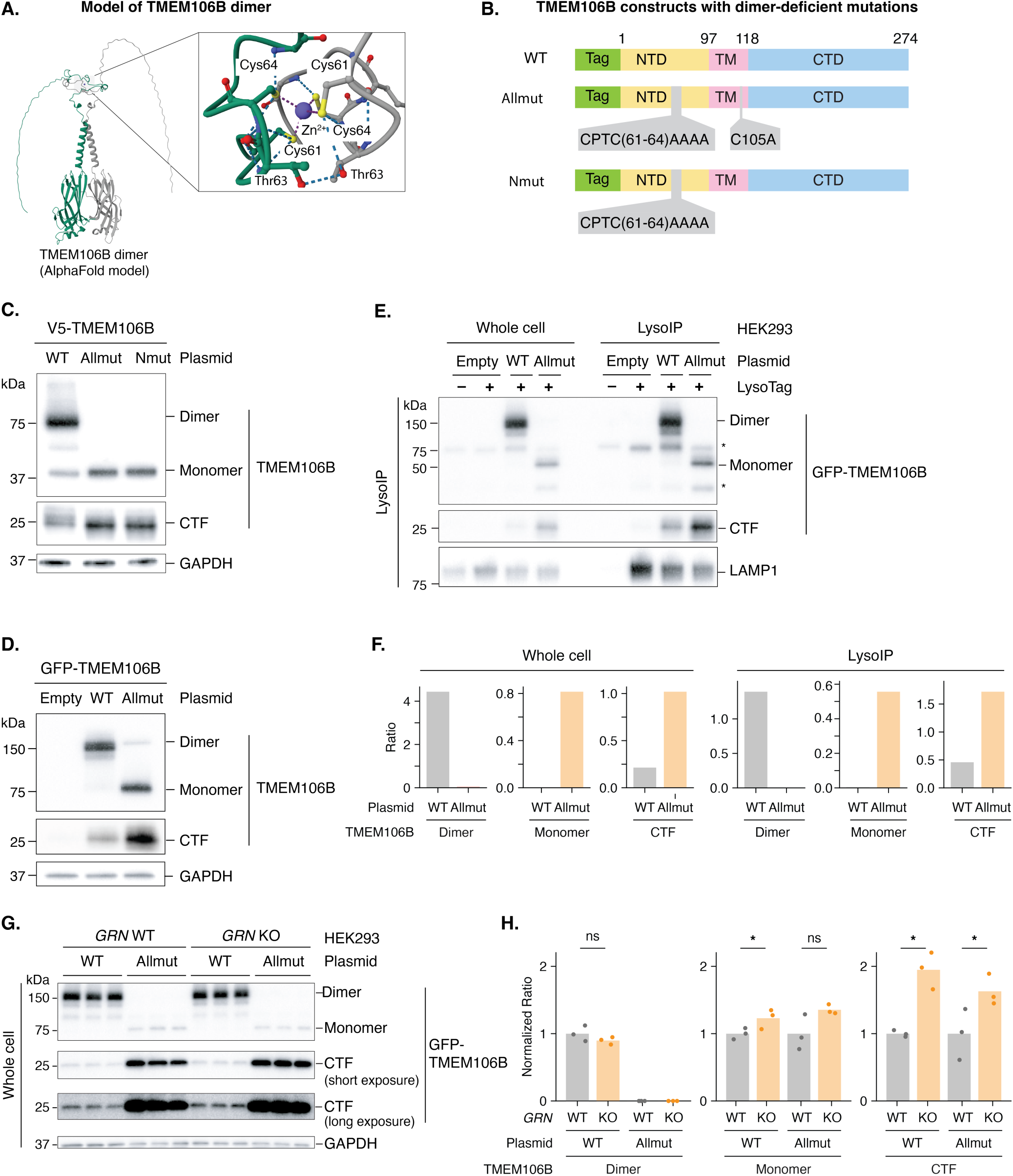
TMEM106B dimer protects against C-terminal fragment accumulation. A. AlphaFold model of TMEM106B homodimer, with a conserved dimer interface mediated by zinc coordination (inset). B. Schematic view of TMEM106B plasmid constructs containing predicted dimer-deficient mutations. (Tag, N-terminal tag; NTD, N-terminal domain; TM, transmembrane; CTD, C-terminal domain). C. Western blots show that mutating predicted zinc-coordinating residues disrupts TMEM106B dimer formation in V5-tagged TMEM106B proteins, leading to the accumulation of the cleaved C-terminal fragment (CTF). D. Western blots show that mutating predicted zinc-coordinating residues disrupts TMEM106B dimer formation in GFP-tagged TMEM106B proteins. E. Western blots show that disrupting TMEM106B dimer formation increases TMEM106B CTF levels in lysosomes. Lysosomes were purified by immunoprecipitation using the LysoTag. Asterisks indicate endogenous TMEM106B dimers and monomers. F. Quantification of TMEM106B dimers, monomers, and CTFs from panel E. Ratios were calculated by dividing the intensity of each TMEM106B species (dimer, monomer, or CTF) by the loading control (LAMP1). G. Western blots show that GRN loss increases C-terminal fragment levels for both WT and Allmut TMEM106B proteins. Short and long exposures of the CTF are shown. H. Quantification of TMEM106B dimers, monomers, and CTFs from panel G. Normalized ratios were calculated by dividing the intensity of each TMEM106B species (dimer, monomer, or CTF) by the loading control (GAPDH) and then normalizing to the values in *GRN* WT condition. Bar plots represent the mean, and each dot represents a replicate (n = 3 replicates per condition). Statistical significance was determined by a one-sided Welch’s t-test: ns (not significant), p > 0.05; *, p ≤ 0.05; **, p ≤ 0.01; ***, p ≤ 0.001; ****, p ≤ 0.0001.

With this TMEM106B C-terminal fragment accumulating mutant in hand, we could now directly test the level at which granulin deficiency acts on TMEM106B. We transfected granulin-deficient (*GRN* KO) and isogenic WT HEK293 cells with plasmids encoding either WT or Allmut *TMEM106B* and performed LysoIP. If granulin regulates TMEM106B dimerization, granulin deficiency would not increase Allmut TMEM106B C-terminal fragments, since Allmut cannot form dimers. Conversely, if granulin acts on C-terminal fragments themselves, granulin deficiency would increase C-terminal fragments for both WT and Allmut TMEM106B. *GRN* deficiency significantly increased C-terminal fragments derived from both WT and Allmut TMEM106B (Figure 2G, H), providing evidence that granulin is not responsible for TMEM106B dimerization but instead acts to reduce cleaved C-terminal fragments.

Does granulin actively promote lysosomal clearance of TMEM106B C-terminal fragments, and is their accumulation upon granulin loss reversible? We performed rescue experiments by supplementing progranulin deficient human neurons with recombinant progranulin protein or bovine serum albumin (BSA), as a negative control. Recombinant progranulin, but not BSA, resulted in a dose-dependent decrease in TMEM106B C-terminal fragment accumulation in progranulin deficient lysosomes (Figure 3A-D). These effects occurred without changes in the levels of full-length TMEM106B (Figure 3A, B), suggesting that granulin specifically modulates lysosomal proteolysis and/or stability of the TMEM106B C-terminal fragment. Thus, granulin deficiency directly causes TMEM106B C-terminal fragment accumulation in lysosomes, likely by impairing the clearance of cleaved C-terminal fragments.

**Figure 3.**
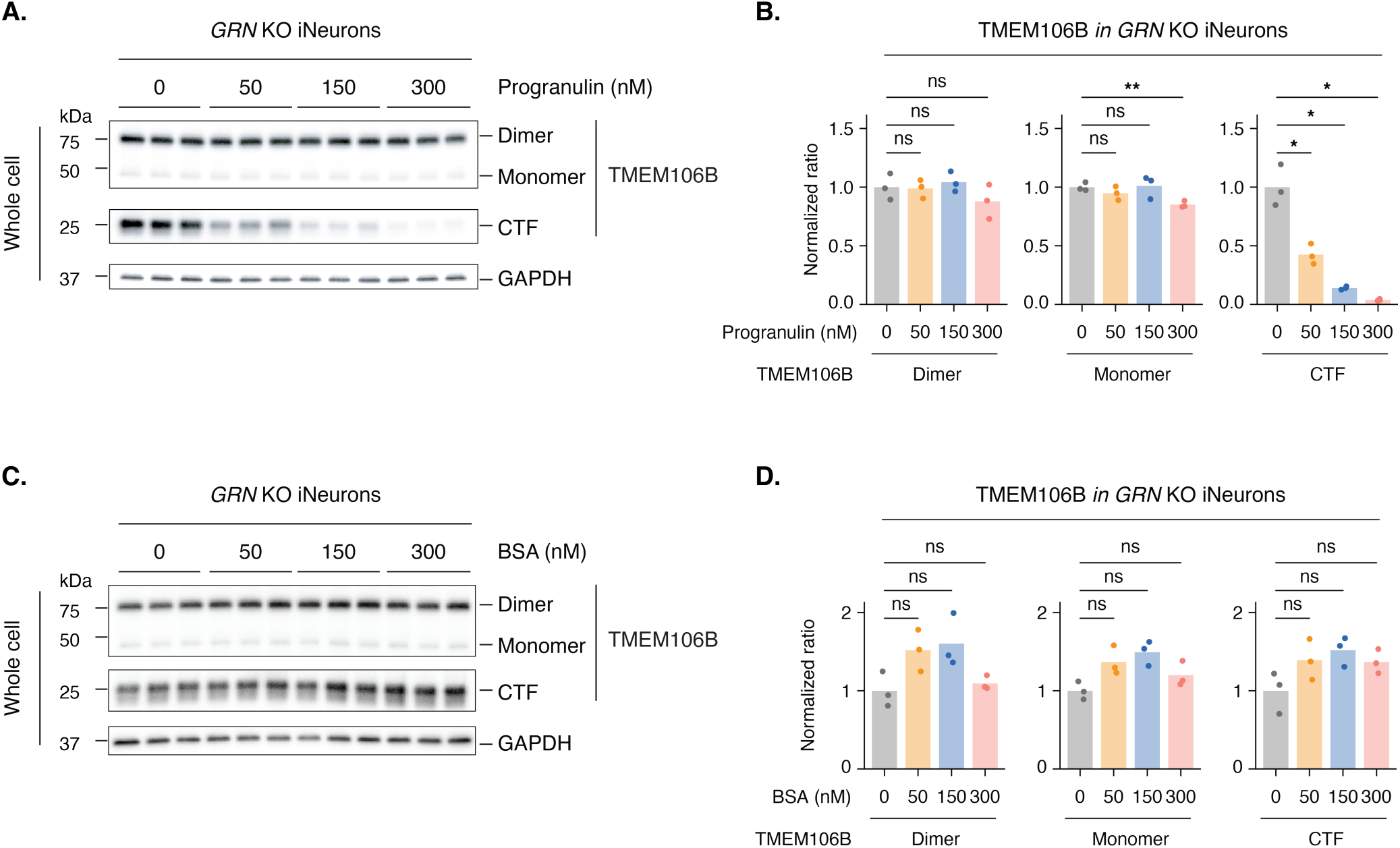
Supplementing progranulin reduces TMEM106B C-terminal fragment accumulation. A. Western blots show that recombinant progranulin reduces TMEM106B CTF accumulation in *GRN* KO iNeurons in a dose-dependent manner. Cells were treated with recombinant progranulin for three days before harvest. B. Quantification of TMEM106B dimers, monomers, and CTFs from panel A. Normalized ratios were calculated by dividing the intensity of each TMEM106B species (dimer, monomer, or CTF) by the loading control (GAPDH), then normalizing to the first bar (0 nM progranulin). C. Western blots show that BSA treatment does not reduce TMEM106B CTF accumulation in *GRN* KO iNeurons in a dose-dependent manner. Cells were treated with BSA for three days before harvest. D. Quantification of TMEM106B dimers, monomers, and CTFs from panel C. Normalized ratios were calculated by dividing the intensity of each TMEM106B species (dimer, monomer, or CTF) by the loading control (GAPDH), then normalizing to the first bar (0 nM BSA). Bar plots represent the mean, and each dot represents a replicate (n = 3 replicates per condition). Statistical significance was determined by a two-sided Welch’s t-test: ns (not significant), p > 0.05; *, p ≤ 0.05; **, p ≤ 0.01; ***, p ≤ 0.001; ****, p ≤ 0.0001.

Because of the striking sensitivity of TMEM106B C-terminal fragment accumulation within lysosomes to granulin deficiency and the potent ability of *TMEM106B* genetic variants to modify risk of FTD in human^13,14,39^, we next asked if and how these two observations might be functionally connected. In other words, do the genetic variants modify FTD penetrance by making TMEM106B either more or less sensitive to lysosomal processing and C-terminal fragment accumulation? But which *TMEM106B* variant to focus on? When *TMEM106B* was initially identified in the FTD genome wide association study (GWAS), there were three significant single nucleotide variants (rs1990622, rs6966915, rs1020004) at the *TMEM106B* locus that were all in high linkage disequilibrium^13^. Later work showed that these variants have a strong impact on age-at-onset in *GRN* mutation carriers with FTD^40^ and that harboring two copies of the protective haplotype conferred protection from autosomal dominant FTD caused by *GRN* mutation^14^. An *Alu* repetitive element located in the *TMEM106B* 3’UTR in linkage disequilibrium with the other variants has also been nominated as contributing to FTD and seems to confer its impact by influencing protein levels^41^. So far, the only *TMEM106B* coding variant linked to the other single nucleotide variants is the missense variant rs3173615, which results in a threonine-to-serine substitution at residue 185. Initial cell-based studies provided evidence that this amino acid change affected TMEM106B protein stability^15^ but there were conflicting reports when this variant was analyzed *in vivo* in *Grn*^−/−^ mice^42^. The threonine variant increases FTD risk whereas the serine variant is protective, reducing or in some cases completely eliminating FTD risk in *GRN* mutation carriers^29^. To define the mechanism, we explored how these variants impact levels of the C-terminal fragment in the lysosome and if and how they intersect with granulin deficiency.

To isolate the effects of the threonine (T) versus serine (S) variant at TMEM106B position 185, we obtained human iPSC lines expressing either the threonine or serine allele on an isogenic background (Figure 4A). We then introduced the LysoTag^33^ to these cell lines, differentiated them into iNeurons, and performed LysoIP to specifically isolate intact lysosomes (Figure S2A). We found a striking risk allele-dose dependent increase in TMEM106B C-terminal fragment accumulation in lysosomes (TT=high; TS=medium; SS=low) (Figure 4B, C; Figure S2B, C). This suggests that the protective effect of the TMEM106B serine variant may lie in preventing pathological lysosomal accumulation (also see Discussion).

**Figure 4.**
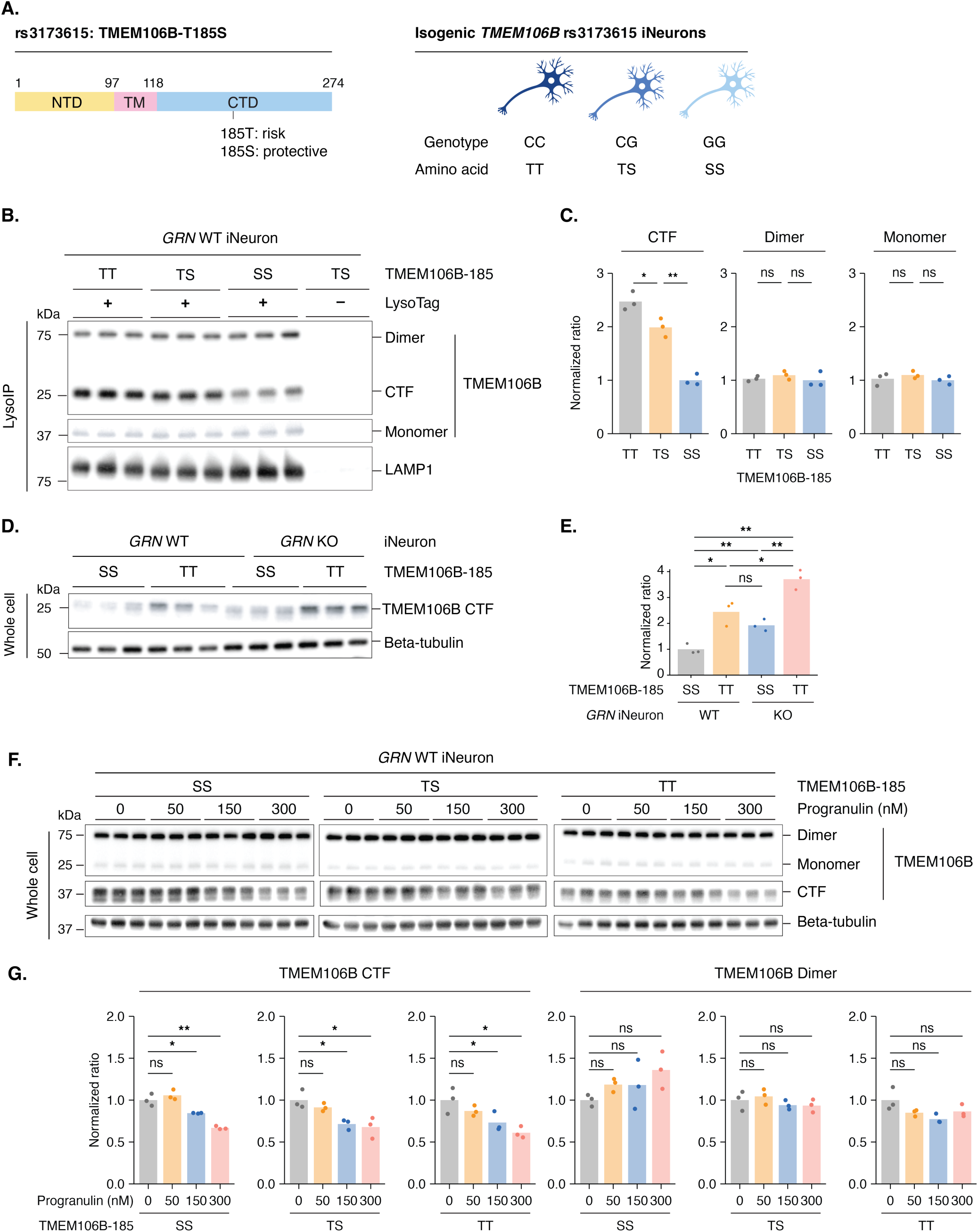
TMEM106B protective coding variant reduces C-terminal fragment accumulation in the lysosome. A. Left panel: Schematic of TMEM106B showing the T185S coding variant (rs3173615) located in the C-terminal domain. Right panel: Isogenic iPSC-derived neurons were generated with three genotypes: CC (homozygous threonine, TT), CG (heterozygous threonine/serine, TS), and GG (homozygous serine, SS). B. Western blots show that the copy number of the protective S185 allele anti-correlates with TMEM106B CTF levels in the lysosome. Purified lysosomes were from *GRN* WT iNeurons with TT, TS, or SS genotypes. Lysosomes were purified by immunoprecipitation using the LysoTag. C. Quantification of TMEM106B dimers, monomers, and CTFs from panel B. Normalized ratios were calculated by dividing the intensity of each TMEM106B species (dimer, monomer, or CTF) by the loading control (LAMP1), then normalizing to the first bar (TT genotype) of each TMEM106B species. D. Western blots show that *GRN* KO increases TMEM106B CTF levels in iNeurons with SS or TT genotypes. Whole cell lysates were analyzed. E. Quantification of TMEM106B CTFs from panel D. Normalized ratios were calculated by dividing the intensity of TMEM106B CTF by the loading control (beta-tubulin) and then normalizing to the first bar. F. Western blots show that recombinant progranulin treatment reduces TMEM106B CTF accumulation in iNeurons with SS, TS, or TT genotypes in a dose-dependent manner. Cells were treated with recombinant progranulin for three days before harvest. G. Quantification of TMEM106B dimers and CTFs from panel F. Normalized ratios were calculated by dividing the intensity of each TMEM106B species (dimer or CTF) by the loading control (beta-tubulin), then normalizing to the first bar (0 nM progranulin) of each genotype group. Bar plots represent the mean, and each dot represents a replicate (n = 3 replicates per condition). Statistical significance was determined by two-sided Welch’s t-test: ns (not significant), p > 0.05; *, p ≤ 0.05; **, p ≤ 0.01; ***, p ≤ 0.001; ****, p ≤ 0.0001.

To further validate these findings and place TMEM106B either upstream or downstream of granulin, we examined TMEM106B C-terminal fragment levels in a granulin knockout context. Remarkably, in the absence of granulin, homozygosity for the protective serine variant reduced the accumulation of the TMEM106B C-terminal fragment in the lysosome to levels comparable to those of the threonine variant in WT cells (Figure 4D, E). In other words, the serine variant seems to blunt the impact of granulin deficiency on TMEM106B C-terminal fragment accumulation in the lysosome. Recombinant progranulin treatment normalized these levels across all genotypes (Figure 4F, G). TMEM106B full-length protein levels remained constant (Figure 4G), indicating that allele-specific effects reflect altered lysosomal processing rather than expression differences. These results provide evidence that the protective effect of the serine variant acts downstream of granulin deficiency.

To extend our findings and examine whether *TMEM106B* protective haplotypes reduce TMEM106B C-terminal fragment levels independent of granulin protein levels *in vivo*, as observed in our cellular models (Figure 4B, C), we analyzed large quantitative proteomic datasets of human brain samples with matched genotype information from the Religious Orders Study and Memory and Aging Project (ROSMAP)^43,44^. We used the first cohort for the primary analysis (ROSMAP-R1, n=358) and the second cohort for confirmatory analysis (ROSMAP-R2; n=185). Inspired by a recent quantitative mass spectrometry study showing that TMEM106B C-terminal domain (CTD)-mapping peptides serve as effective indicators of C-terminal fragment accumulation^45^, we examined whether *TMEM106B* rs3173615 and *GRN* rs5848 variants independently affect these peptide levels.

Consistent with our observations in TMEM106B-185 isogenic cell lines (Figures 4B, C), the *TMEM106B* protective allele (rs3173615-G, which encodes a serine residue at position 185 of the protein instead of threonine) was significantly associated with a robust, dose-dependent reduction of CTD-mapping peptide levels (Figure 5A; Figure S3B) and modestly associated with increased levels of a few NTD-mapping peptides (Figure 5B; Figure S3C) in both cohorts. Furthermore, a common *GRN* variant (rs5848-T) moderately reduces granulin protein levels^46^. Indeed, we found a dose-dependent reduction with T allele copy number in ROSMAP data (Figure S3A). Remarkably, rs5848-T was also significantly associated with increased TMEM106B CTD-mapping peptides but not NTD-mapping peptides in the first cohort (Figure 5C, D; Figure S3D, E). These results directly support our finding that granulin deficiency drives lysosomal TMEM106B C-terminal fragment accumulation (Figure 1A, B).

**Figure 5.**
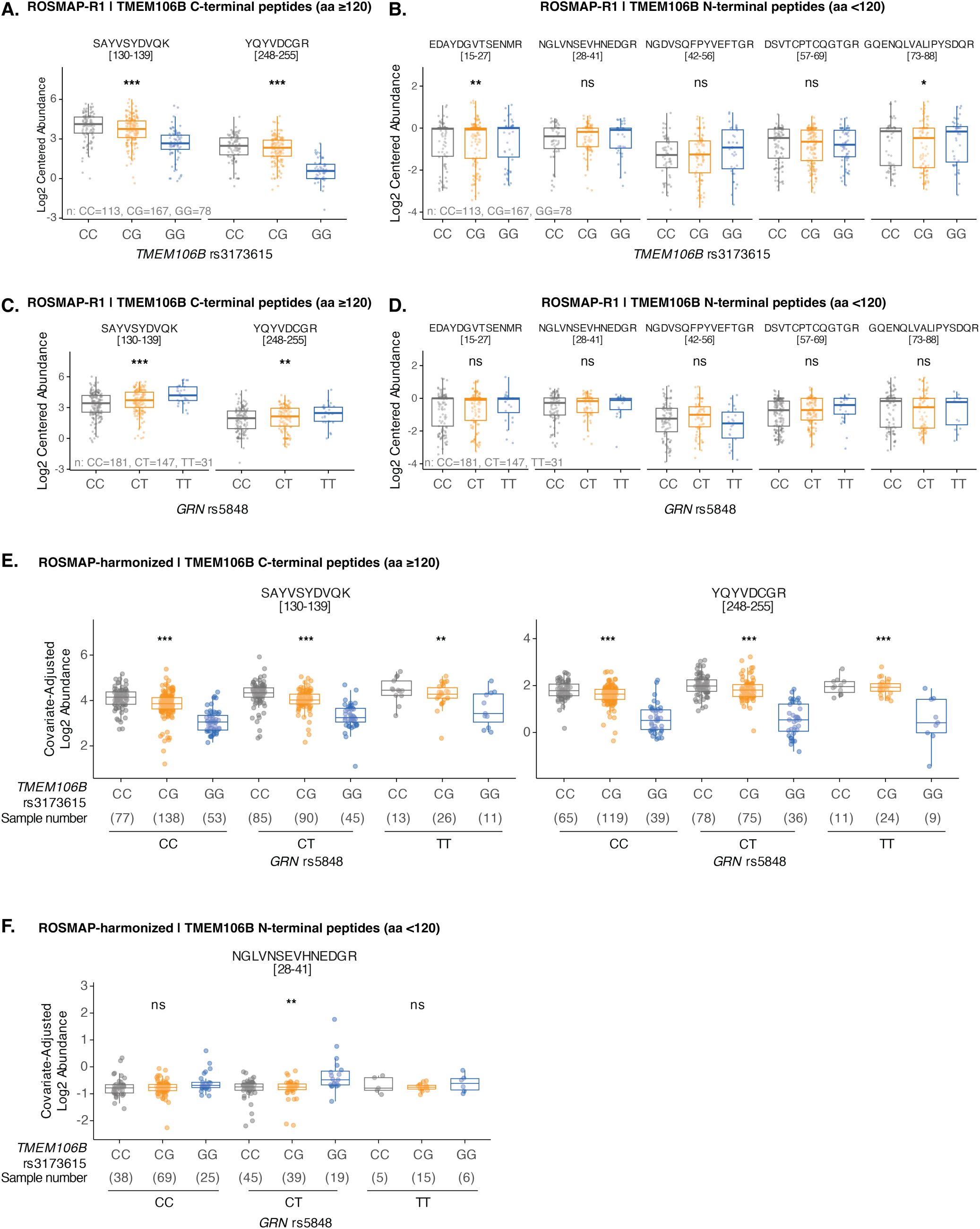
The TMEM106B risk haplotype and common GRN variant independently drive TMEM106B C-terminal fragment accumulation in human brain. A. Box plots showing that the protective *TMEM106B* rs3173615 allele is significantly associated with reduced levels of peptides mapping to the TMEM106B C-terminal domain (CTD) in the first ROSMAP cohort (ROSMAP-R1). Association analysis was performed using a linear regression model adjusted for sample batch, sex, race, age of death, postmortem interval, and *APOE* genotype. B. Box plots showing that the protective *TMEM106B* rs3173615 allele is associated with increased levels of select peptides mapping to the TMEM106B N-terminal domain (NTD) in the first ROSMAP cohort (ROSMAP-R1). The association analysis was performed as in panel A. C. Box plots showing that the common *GRN* risk variant (rs5848-T) is significantly associated with increased levels of TMEM106B CTD-mapping peptides in the first ROSMAP cohort (ROSMAP-R1). The association analysis was performed as in panel A. D. Box plots showing that the common *GRN* variant (rs5848-T) is not significantly associated with levels of TMEM106B NTD-mapping peptides in the first ROSMAP cohort (ROSMAP-R1). The association analysis was performed as in panel A. E. Covariate-adjusted box plots illustrating that the protective *TMEM106B* rs3173615 allele is associated with a stepwise reduction of CTD-mapping peptides across all *GRN* genotypes in the harmonized ROSMAP data, displaying the independent, additive effects of both variants. A covariate-adjusted linear model was used to estimate adjusted *p* values for the *TMEM106B* simple effect within each *GRN* genotype stratum, and resulting adjusted p values are indicated. F. Covariate-adjusted box plots showing the N-terminal TMEM106B peptide across GRN genotypes in the harmonized ROSMAP data. The *TMEM106B* protective rs3173615 allele is significantly associated with increased N-terminal peptide levels in the additive joint model, whereas GRN was not. Adjusted *p* values for the *TMEM106B* simple effect within each *GRN* genotype stratum were calculated as in panel E. Details of statistical analysis can be found in the Methods section. Statistical significance labels are ns (not significant), p > 0.05; *, p ≤ 0.05; **, p ≤ 0.01; ***, p ≤ 0.001; ****, p ≤ 0.0001.

To look for possible genetic interactions between these *TMEM106B* and *GRN* alleles, we used both continuous linear modeling and categorical interaction (ANOVA) framework-based modeling, adjusting for batch, sex, age of death, and postmortem interval covariates. We did not observe a statistically significant genetic interaction between *TMEM106B* rs3173615 and *GRN* rs5848 alleles on TMEM106B peptide abundances in these datasets (adjusted *p* value > 0.05). In contrast, the joint additive model without an interaction term showed independent associations of both loci with the two C-terminal TMEM106B peptides: SAYVSYDVQK and YQYVDCGR (*TMEM106B* adjusted *p* value = 9.98×10^−39^ and 7.95×10^−46^; *GRN* adjusted *p* value = 5.28×10^−8^ and 0.0024, respectively). Importantly, when visualizing covariate-adjusted abundances stratified by *GRN* rs5848 genotype, the protective *TMEM106B* rs3173615 allele was significantly associated with a stepwise reduction of CTD-mapping peptides but not NTD-mapping peptides across all *GRN* genotypes (Figure 5E, F; Figure S3F, G). In other words, in human brain, the presence of the *TMEM106B* protective allele affords dose-dependent protection against granulin deficiency-associated TMEM106B C-terminal fragment accumulation. This observation provides an explanation for why homozygous *TMEM106B* protective allele carriers have near lifetime resilience to granulin deficiency. Together, these findings corroborate our *in vitro* findings that the TMEM106B protective mechanism effectively promotes TMEM106B C-terminal fragment clearance independent of *GRN* haploinsufficiency, providing a mechanistic connection between *TMEM106B*, the strongest genetic risk factor, and granulin deficiency in FTD. Taken together, we propose that granulin deficiency leads to the accumulation of C-terminal fragments in lysosomes and that the *TMEM106B* protective allele reduces C-terminal fragment stability, preventing its accumulation and subsequent amyloid formation.

## Discussion

These findings, and similar ones by Ward, Mosalaganti, Petrucelli, and colleagues in an accompanying manuscript, unify *GRN* and *TMEM106B* within a single lysosomal proteostasis pathway that determines FTD risk. Granulin deficiency promotes accumulation of TMEM106B C-terminal fragments in the lysosome, a process mitigated by granulin supplementation and by protective *TMEM106B* alleles. Previous work has shown a *TMEM106B* risk haplotype association between increased TMEM106B C-terminal fragments and reduced dimers in human brain^32,47^ but our findings establish granulin as a direct regulator of TMEM106B C-terminal fragment clearance and reveal that lysosomal TMEM106B C-terminal fragment formation is not merely a correlate of aging or disease but instead a key player in the pathogenic process. This pathway also explains why *GRN* mutation carriers homozygous for the *TMEM106B* protective allele remain clinically unaffected even in the face of granulin deficiency^14^.

Our structural and genetic experiments identify TMEM106B dimerization as a critical determinant of lysosomal stability. Disruption of the dimer interface enhances C-terminal fragment accumulation. Since the S or T residue at position 185 of TMEM106B resides within the N-X-S/T consensus sequence for N183 glycosylation^22^ and N-X-S typically shows lower glycosylation occupancy than N-X-T^48^, TMEM106B with the protective S185 allele might be less efficiently glycosylated at N183, thereby promoting clearance of the C-terminal fragment. Importantly, the TMEM106B protective S185 allele lowered C-terminal fragment levels even in the granulin deficiency background (Figures 4 and 5), which provides a molecular explanation for the near-complete protection observed in human genetics and aligns with recent structural observations of TMEM106B fibrils in aged brains^24–27^ and the accumulation of TMEM106B C-terminal fragments in FTD patient brain in a risk haplotype-dependent manner^32^. Together, these results reveal how genetic variation shapes lysosomal resilience in neurodegeneration. Therapeutically, enhancing granulin levels^49–52^ or stabilizing TMEM106B dimers may restore lysosomal proteostasis (Figure 2) and prevent disease. More broadly, this work exemplifies how protective alleles can reveal intrinsic buffering mechanisms against protein aggregation, guiding strategies to strengthen cellular defense systems in neurodegenerative disorders.

## Acknowledgments

We acknowledge funding support from the following sources: postdoctoral scholar awards from the Phil and Penny Knight Initiative for Brain Resilience, Stanford University (Y.Z. and J.X.); a grant from the Larry L. Hillblom Foundation (Y.Z.); a Target ALS Springboard Fellowship (Y.Z.); a Live Like Lou Graduate Fellowship (T.P.N.); a Stanford Graduate Fellowship (T.P.N); the NIH (R35NS137159, U54NS123743, R01AG064690), an Alzheimer’s Disease Strategic Fund grant, and Target ALS to A.D.G. A.D.G. is a Biohub – San Francisco Investigator. This work was partially supported by grants from Beatbatten, the NCL Foundation (NCL-Stiftung), the Knight Initiative for Brain Resilience at Stanford University, Arc Institute, and the Chan Zuckerberg Initiative Neurodegeneration Challenge Network (2024-338545) to M.A-R. M.A-R is a Stanford Terman Fellow, a Sloan Fellow, Arc Institute Innovation Investigator, and a Pew-Stewart Scholar for Cancer Research, supported by The Pew Charitable Trusts and The Alexander and Margaret Stewart Trust. We would like to thank Leonard Petrucelli (University of Miami) for sharing the TMEM106B C-terminal antibody, Michael Ward (NIH) for sharing TMEM106B isogenic iPSC lines, and Hua Tang (Stanford) for her suggestions on genotype-peptide analysis. Some of the computing for this project was performed on the Sherlock cluster. We would like to thank Stanford University and the Stanford Research Computing Center for providing computational resources and support that contributed to these research results.

The results published here are in part based on data obtained from The AD Knowledge Portal (https://doi.org/10.7303/9618239). Study data were provided through the Accelerating Medicine Partnership for AD (U01AG046161 and U01AG061357) based on samples provided by the Rush Alzheimer’s Disease Center, Rush University Medical Center, Chicago. Data collection was supported through funding by NIA grants P30AG10161, R01AG15819, R01AG17917, R01AG30146, R01AG36836, U01AG32984, U01AG46152, the Illinois Department of Public Health, and the Translational Genomics Research Institute.

## Supplementary figures

**Figure S1.**
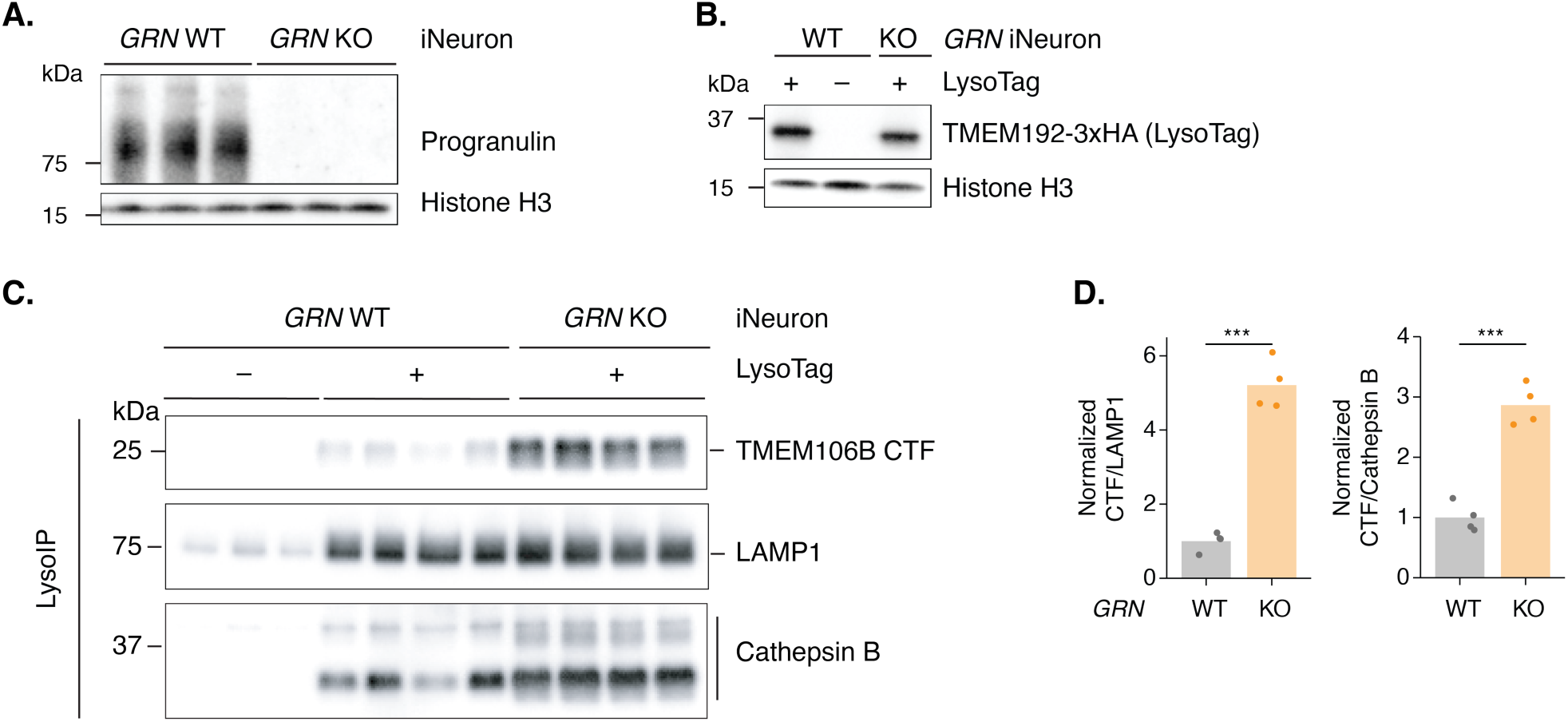
GRN loss increases levels of TMEM106B cleaved C-terminal fragments in the lysosome. A. Western blot confirms that *GRN* is knocked out in the *GRN* KO cell line. B. Western blot confirms robust expression of the LysoTag (TMEM192-3xHA) in iNeurons. C. Western blots show that *GRN* knockout (KO) in iNeurons leads to accumulation of TMEM106B C-terminal fragments (CTFs) in lysosomes. Purified lysosomes were from *GRN* wild-type (WT) iNeurons without LysoTag, *GRN* WT iNeurons with LysoTag, and *GRN* KO iNeurons with LysoTag. Samples were from the same experiment as in Figure 1A. D. Quantification of TMEM106B CTFs from panel C. Normalized ratios were calculated by dividing the intensity of the TMEM106B CTF by either LAMP1 or Cathepsin B, then normalizing to the first bar (*GRN* WT with LysoTag for purified lysosomes). Bar plots represent the mean, and each dot represents a replicate (n = 3 replicates per condition). Statistical significance was determined by a two-sided Welch’s t-test: ns (not significant), p > 0.05; *, p ≤ 0.05; **, p ≤ 0.01; ***, p ≤ 0.001; ****, p ≤ 0.0001.

**Figure S2.**
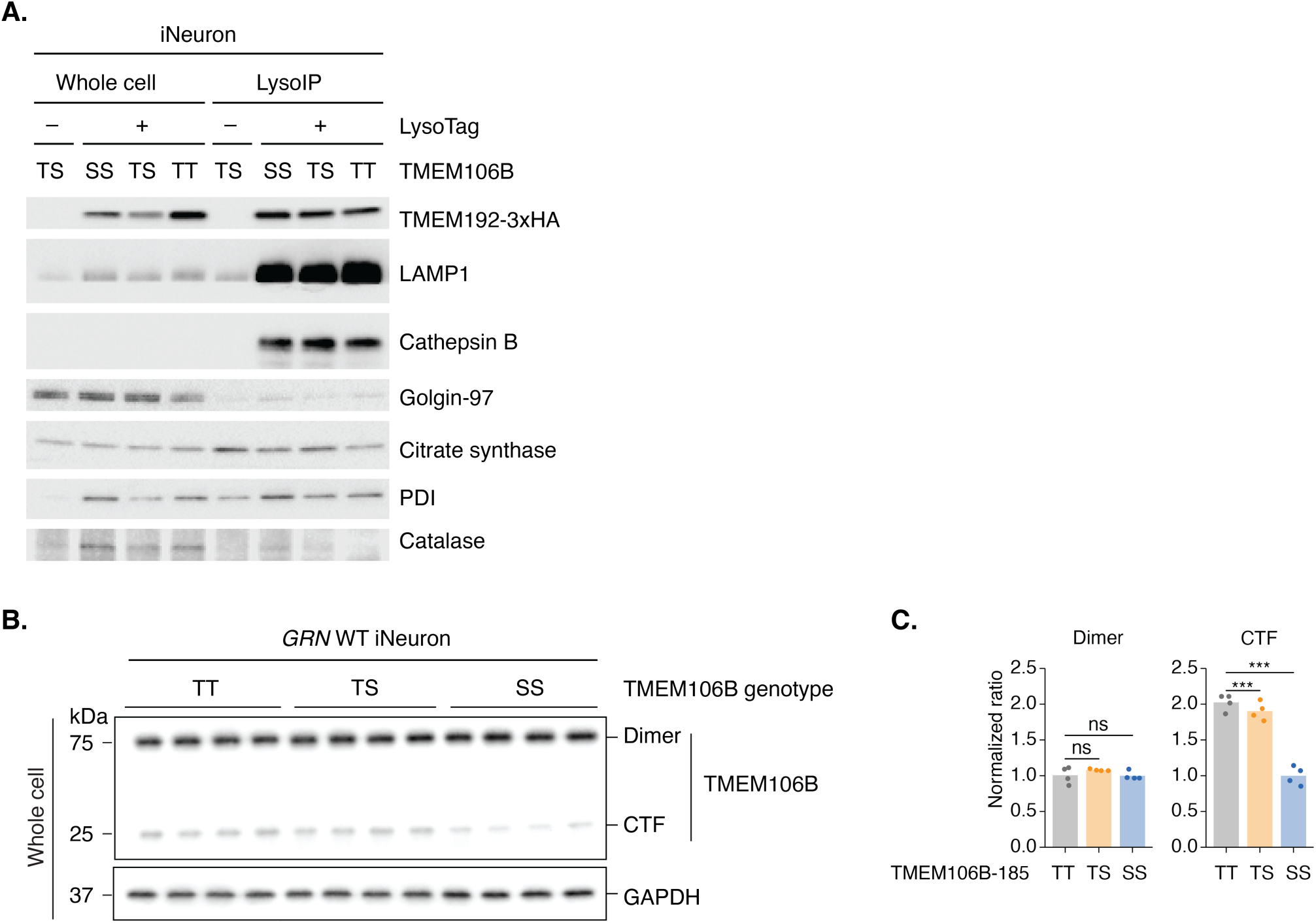
TMEM106B protective alleles are associated with reduced C-terminal fragment accumulation. A. Western blots show that LysoIP in iNeurons specifically purifies intact lysosomes without enriching other organelles. Western blots of whole cell lysates and purified lysosomes from LysoTag-expressing *GRN* WT iNeurons harboring different copy numbers of the TMEM106B protective coding variant (SS, TS, and TT) and *GRN* WT iNeurons harboring a heterozygous TMEM106B coding variant (TS) without a LysoTag as a negative control. Organelle markers are labeled. B. Western blots show that the copy number of the protective S185 allele anti-correlates with TMEM106B CTF levels in iNeurons. Whole cell lysates were from *GRN* WT iNeurons with TT, TS, or SS genotypes. C. Quantification of TMEM106B dimers and CTFs from panel B. Normalized ratios were calculated by dividing the intensity of each TMEM106B species (dimer or CTF) by the loading control (GAPDH), then normalizing to the first bar (TT genotype) of each TMEM106B species. Bar plots represent the mean, and each dot represents a replicate (n =3 replicates per condition). Statistical significance was determined by two-sided Welch’s t-test: ns (not significant), p > 0.05; *, p ≤ 0.05; **, p ≤ 0.01; ***, p ≤ 0.001; ****, p ≤ 0.0001.

**Figure S3.**
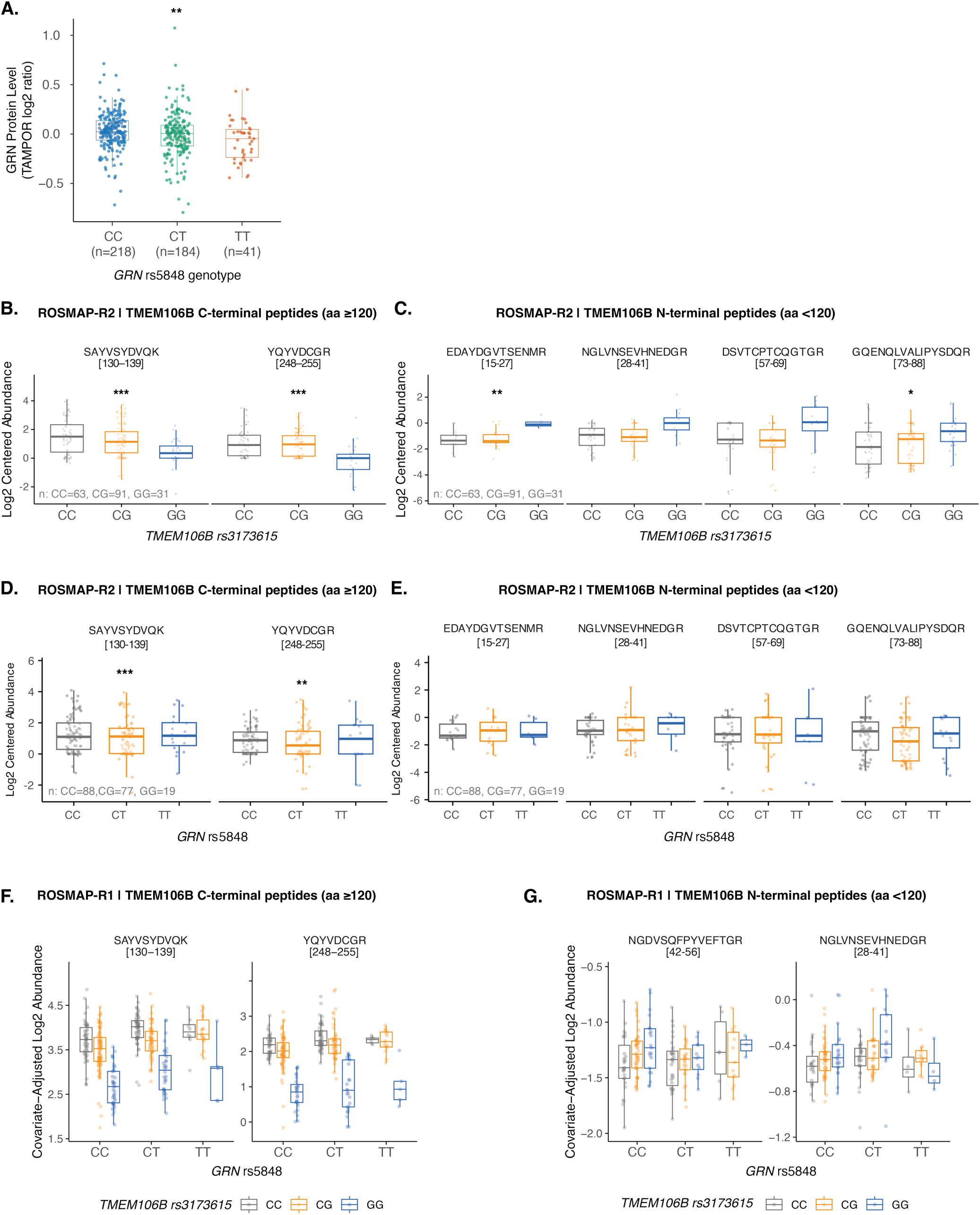
The TMEM106B risk haplotype and common GRN variant independently drive TMEM106B C-terminal fragment accumulation in human brain. A. Box plots showing that the common *GRN* risk variant (rs5848-T) is significantly associated with reduced progranulin protein levels. The association analysis was performed as using limma, adjusting for sex, postmortem interval, and final consensus cognitive diagnosis. B. Box plots showing that the protective *TMEM106B* rs3173615 allele is significantly associated with reduced levels of peptides mapping to the TMEM106B C-terminal domain (CTD) in the second ROSMAP cohort (ROSMAP-R2). Association analysis was performed using a linear regression model adjusted for sample batch, sex, race, age of death, postmortem interval, and *APOE* genotype. C. Box plots showing that the protective *TMEM106B* rs3173615 allele is significantly associated with increased levels of peptides mapping to the TMEM106B N-terminal domain (NTD) in the second ROSMAP cohort (ROSMAP-R2). The association analysis was performed as in panel B. D. Box plots showing that the common *GRN* risk variant (rs5848-T) is not significantly associated with increased levels of TMEM106B CTD-mapping peptides in the second ROSMAP cohort (ROSMAP-R2). The association analysis was performed as in panel B. E. Box plots showing that the common *GRN* variant (rs5848-T) is not significantly associated with levels of TMEM106B NTD-mapping peptides in the second ROSMAP cohort (ROSMAP-R2). The association analysis was performed as in panel B. F. Covariate-adjusted box plots illustrating that the protective *TMEM106B* rs3173615 allele is associated with a stepwise reduction of CTD-mapping peptides across all *GRN* genotypes in in the first ROSMAP cohort (ROSMAP-R1), displaying the independent, additive effects of both variants. G. Covariate-adjusted box plots showing N-terminal TMEM106B peptides across GRN genotypes in harmonized ROSMAP data. Details of the statistical analysis can be found in the Methods section. Statistical significance was determined by two-sided Welch’s t-test: ns (not significant), p > 0.05; *, p ≤ 0.05; **, p ≤ 0.01; ***, p ≤ 0.001; ****, p ≤ 0.0001.

## Materials and Methods

### Stem cell maintenance and differentiation into iNeurons

The following human induced pluripotent stem cells (iPSCs; KOLF2.1J) were purchased from The Jackson Laboratory: KOLF2.1J, *GRN* knockout, TMEM106B-185SS, TMEM106B-185TS, and TMEM106B-185TT lines. These iPSC lines were maintained in mTeSR Plus media (StemCell Technologies, 100-0276) with 50 U/mL Penicillin-Streptomycin (Gibco, 15140122) on plates coated with Matrigel (Corning, 354230). Cells were fed every two days and split every 3-4 days using ReLeSR (StemCell Technologies, 100-0483) according to the manufacturer’s instructions. Cells were split using mTeSR Plus with 10 nM of ROCK1 inhibitor (Tocris, 1254), and media was changed to media without ROCK1 inhibitor within 24 hours. The differentiation of iPSCs to neurons by forcing *NGN2* overexpression was carried out as previously described^53^. In brief, cells were transduced with a Tet-On induction system to drive the expression of the transcription factor *NGN2*. Cells were dissociated on day 3 of differentiation and replated on poly-D-lysine and laminin double-coated tissue culture plates in Neurobasal Medium (Thermo Fisher, 21103049) containing neurotrophic factors, BDNF (Peprotech, 450-10) and GDNF (Peprotech, 450-03).

### Generation of LysoTag expressing stable cell lines

To generate stable cell lines expressing LysoTag (TMEM192-3×HA) for lysosome purification, the PiggyBac (PB) transposon system was utilized. Human iPSCs were seeded and maintained until reaching 70–80% confluence in mTeSR Plus media (StemCell Technologies, 100-0276) with 50 U/mL Penicillin-Streptomycin (Gibco, 15140122) on plates coated with Matrigel (Corning, 354230). For electroporation, 5×10^5^ cells were processed with 1 µg of LysoTag-PB plasmid and 0.5 µg of a PB helper transposase-encoding plasmid. Electroporation was performed using the Lonza 4D Nucleofector X according to the manufacturer’s protocol, followed by immediate plating onto one well of a 24-well plate coated with Matrigel. Media was replaced with fresh medium containing 1 µg/mL puromycin 24 hours post-electroporation to facilitate transposition and genomic integration. Cells were maintained under antibiotic selection for a minimum of seven days prior to expansion into monolayers or cryopreservation. Expression of LysoTag in iPSCs and iNeurons was confirmed by western blot using an HA antibody. The TMEM192-3×HA iPSC lines were used for engineering all subsequent gene deletions.

### Generation of GRN knockout cell lines

To generate *GRN*-deficient lines, CRISPR/Cas9-mediated genome editing was performed in human iPSCs. Two sgRNAs targeting the *GRN* gene locus were designed (GCAAAGTACCAAGGAACGTC and TAAGGCCTTCCCTGTCAGAA), along with a single-stranded DNA (ssDNA) template containing homology arms flanking the Cas9 cleavage sites for homology-directed repair (HDR) (CTGAGTGACCCTAGAATCAAGGGTGGCGTGGGCTTAAGCAGTTGCCAGADHAAGGGGGTTGTGG CAAAAGCCACATTACAAGCTGCCATCCCCTCCCCGT). The ssDNA HDR template contains mixed bases to increase the knock out efficiency (D=A, G, T; H=A, C, T) The assembled CRISPR/Cas9 complex and the ssDNA HDR template were electroporated into 5×10^5^ iPSC cells using the Lonza 4D Nucleofector X System according to the manufacturer’s protocol. Following electroporation, cells were plated at low density in 96-well plates to isolate single clones. Successful *GRN* deletion was validated by PCR screening, Sanger sequencing, and western blot analysis using anti-progranulin antibodies.

### Mice

Mice were acquired and maintained, as previously described^54^. The mice were housed in a controlled environment with regulated 20–26 °C temperature and 30–70% relative humidity, 12-hour light/dark cycles and access to food and water ad libitum. Supplies were checked daily, and the cages were cleaned every 4–5 days. All mouse procedures were conducted in accordance with the approved guidelines of the Administrative Panel on Laboratory Animal Care at Stanford University. *Grn* KO mice (JAX strain no. 013175) were crossed with LysoTag mice (JAX strain no. 035401) to generate LysoTag mice with the desired *Grn* KO genotypes. Mice aged 6 months were used for all experiments.

Mouse genotypes were confirmed by PCR according to the vendor instructions. PCRs were performed with primers: 11080 (common): 5’-AGAGGGTGAGCTGCAATGTT-3’; 11081 (wild-type reverse): 5’-AAGGGCATTAGCCAAGTGTG-3’; 11082 (mutant reverse): 5’-TCTCCCAGGTAGCCCCTACT-3’, using a program of 5 min at 95 °C, followed by 35 cycles of 1 min at 95 °C, 1 min at 55 °C, and 2 min at 72 °C, and finally with a 10-min incubation at 72 °C. The WT allele yields a band of 468 bp, and the mutant allele yields a band of 211 bp.

### Purification of recombinant progranulin

Expi293F cells were cultured using Expi293 media and transfected with the 2XStrep-PGRN plasmids at a density of 3×10^6^ cells per mL. Transfection was performed with PEI Star (Tocris) at a ratio of 5 µL of 1 mg/mL PEI to 1 µg plasmid DNA per milliliter of cells. Following transfection, glucose and valproic acid were added immediately to reach the final concentrations of 0.4% glucose (w/v) and 0.05% valproic acid (w/v). Equivalent amounts of valproic acid and glucose were added 2 days after transfection, and cells were harvested 3 days after transfection. The culture medium was harvested and filtered through a 0.22-µm PES filter to remove cell debris and then applied to Strep-Tactin®XT 4Flow® high-capacity columns (IBA Bioscience). The columns were then washed with 50 mL of wash buffer (100 mM Tris-HCl, 150 mM NaCl, 1 mM EDTA, pH 8). Proteins were then eluted using elution buffer (100 mM Tris-HCl, 150 mM NaCl, 1 mM EDTA, 50 mM biotin, pH 8). Using an Amicon 3-kDa MWCO concentrator, the eluted protein was concentrated and further purified by size exclusion chromatography on a Superdex 200 10/300 column. The purity of each fraction was determined by SDS-PAGE followed by Coomassie staining. Pure fractions were pooled and concentrated.

### Progranulin supplementation

Seven days after differentiation, iNeurons were cultured in media containing the indicated concentrations of recombinant progranulin or BSA for three days before harvest for downstream analysis.

### Computational Modeling of TMEM106B Dimerization

To elucidate the structural determinants of TMEM106B dimerization, homodimeric assembly was modeled using AlphaFold 3. Initial modeling performed without ligand inputs identified a conserved interface resembling the Rad50 zinc hook motif containing CXXC pairs from opposing monomers. A second modeling run incorporating explicit Zn²⁺ ions as ligands validated this configuration, revealing high-confidence sulfur-metal contacts that specifically coordinate an intermolecular zinc ion via Cys61 and Cys64 within each monomer. These residues were therefore inferred to stabilize the dimeric interface through metal coordination.

### Generation of TMEM106B dimer-deficient mutants

To validate the structural predictions regarding dimerization interfaces, site-directed mutagenesis was used to introduce specific amino acid substitutions into the TMEM106B coding sequence. The following mutant constructs were generated: Allmut (CPTC61-64→AAAA; C105A) and Nmut (CPTC61-64→AAAA). To facilitate protein analysis, an N-terminal GFP or V5 tag was fused to all constructs. Final plasmids were verified by sequencing before use in cellular experiments.

### qRT-PCR to examine TMEM106B mRNA levels

Total RNA (350 ng) from *GRN* WT or *GRN* KO iPSCs was extracted and reverse transcribed to cDNA using the PrimeScript™ RT Reagent Kit with gDNA Eraser (Takara, RR047A). qPCR was carried out using the PowerUp™ SYBR™ Green Master Mix kit and detected using the QuantStudio3 system (Thermo Fisher). The following *TMEM106B* primers were used, as previously described^55^: TMEM106B_total_forward (GCCTCCATCATGACACACTTAC) and TMEM106B_total_reverse (GGGGCATTTCTATTTGTTTGCT).

### Lysosome purification using LysoIP

Lysosomes were purified from livers of mice and iNeurons, following established protocols^33,34,54^. The livers of mice were obtained after euthanasia. Small uniform liver samples were collected using a 6-mm-diameter biopsy punch to minimize variability between samples. Collected liver tissues were homogenized in PBS containing protease inhibitors with a Dounce homogenizer immediately with 25 strokes. The resulting lysates were centrifuged at 1,000×g at 4 °C for 2 min. The supernatant was then transferred to a 1.6 mL microcentrifuge tube containing 10 µL of anti-hemagglutinin magnetic beads and incubated at 4 °C for 15 min with rotational mixing. For iNeurons, one 15 cm dish of iNeurons was collected and resuspended in 1-mL of PBS containing protease inhibitor and processed the same as liver samples.

### Immunoblotting

Cells were washed with ice-cold PBS once and lysed at 4 °C for 15 min in ice-cold RIPA buffer (Sigma-Aldrich, R0278) supplemented with a protease inhibitor cocktail (Thermo Fisher Scientific, 78429) and a phosphatase inhibitor cocktail (Thermo Fisher Scientific, 78426). Lysed cells were then centrifuged at 20,000×g for 15 min at 4 °C, and the supernatant was collected. Protein concentrations were determined using bicinchoninic acid assays (Invitrogen, 23225). Protein lysates were then normalized and mixed with 4× Laemmli buffer (Bio-Rad, 1610747) without including beta-mercaptoethanol. For protein lysates from cells, 10 µg of protein lysates were used for each sample. For protein lysates from LysoIP experiments, 10 µL of lysates were used for each sample. Lysates were loaded onto 10-20% Tris-Glycine mini gels (Thermo Fisher Scientific, XP10200BOX) for gel electrophoresis on ice at 100 volts for 2 hours and transferred onto 0.45-μm polyvinylidene difluoride membranes (Bio-Rad, 162-0115) at 250 mA for 2 hours using the wet transfer method (Bio-Rad Mini Trans-Blot Electrophoretic Cell, 170-3930). Membranes were blocked in EveryBlot Blocking Buffer (Bio-Rad, 12010020) or 5% non-fat dry milk in 1× TBST for 1 hour and then incubated overnight at 4 °C in blocking buffer containing the primary antibodies. Membranes were subsequently incubated in blocking buffer containing secondary antibodies for 1 hour. Blots were imaged via the ChemiDoc XRS+ System (Bio-Rad) using SuperSignal West Femto Maximum Sensitivity Substrate (Thermo Fisher Scientific, 34094), or via the LI-COR Odyssey CLx Imager. The intensity of bands was quantified using Fiji and normalized to the corresponding controls.

Primary antibodies used in this work with dilution information are as follows: TMEM106B (E7H7Z) antibody (1:500; Cell Signaling Technology, 93334), cleaved TMEM106B (Ser120) antibody (1:500; Cell Signaling Technology, 87145), C-terminal TMEM106B antibody (1:1000, created in the Dr. Leonard Petrucelli laboratory), GAPDH (1:2000; Sigma-Aldrich, G8795), Histone H3 antibody (1:5000; Abcam, ab1791), Beta-tubulin antibody (1:40000, Sigma-Aldrich, 66240-1-Ig), Human LAMP1 antibody (1:1000, Cell Signaling Technology, 9091P or 15665S), Mouse LAMP1 antibody (1:1000, DSHB, 1D4B), progranulin antibody (1:1000, R&D Systems, AF2420), CTS B (1:1000, Cell Signaling Technology, 31718T), PDI antibody (1:1000, Enzo Life Sciences, ADI-SPA-891-D), Citrate synthase antibody (1:1000, Cell Signaling Technology, 14309T), Golgin-97 antibody (1:1000, Cell Signaling Technology, 13192T), HA-Tag (C29F4) antibody (1:1000, Cell Signaling Technology, 3724S), Catalase antibody (1:1000, Cell Signaling Technology, D4P7B), GFP antibody (1:2000, Antibodies Incorporated, 75-131), and V5 antibody (1:1000, Thermo Fisher Scientific, R960-25).

Secondary antibodies used in this work with 1:5000 dilution are as follows: Horseradish peroxidase-conjugated anti-rabbit IgG (H + L; Life Technologies, 31462), horseradish peroxidase-conjugated anti-mouse IgG (H + L; Thermo Fisher Scientific, 62-6520), horseradish peroxidase-conjugated anti-rat IgG (H + L; Thermo Fisher Scientific, 31470), horseradish peroxidase-conjugated anti-goat IgG (H + L; Santa Cruz Biotechnology, sc-2313), IRDye 800CW anti-rabbit IgG (H+L; LicorBio, 926-32211), IRDye 800CW anti-mouse IgG (H+L; LicorBio, 926-32210), IRDye 800CW anti-goat IgG (H+L; LicorBio, 926-32214), IRDye 800CW anti-rat IgG (H+L; LicorBio, 926-32219), and IRDye 680CW anti-mouse IgG (H+L; LicorBio, 926-68070).

### Progranulin protein level analysis by rs5848 genotype (ROSMAP TMT proteomics)

Dorsolateral prefrontal cortex TMT quantitative proteomics data were obtained from the Religious Orders Study and Memory and Aging Project (ROSMAP) Study (syn17015098). Two TMT experiments were processed: Batch 1 (50 TMT batches, 10-plex; n = 400 participants) and Batch 2 (14 TMT batches, 16-plex; n = 210 participants). Proteins with >50% missing values were excluded. Within-batch normalization was performed using TAMPOR^56^, which computes log2 abundance ratios against a batch-specific global internal standard channel. The 7,502 proteins detected in both batches were retained for cross-batch analysis. Batch harmonization was performed using ComBat, yielding a combined matrix of 7,502 proteins × 604 unique participants.

*GRN* rs5848 genotypes were extracted from ROSMAP whole-genome sequencing data (hg19) and mapped to participant IDs using ROSMAP metadata. Genotypes were encoded as T-allele dosage (CC = 0, CT = 1, TT = 2). Progranulin (GRN; UniProt P28799) protein levels were tested for association with rs5848 dosage using a linear model adjusted for sex, post-mortem interval, and cognitive diagnosis (limma^57^) in N = 518 participants. A proteome-wide scan of all 7,502 proteins was additionally performed, with FDR correction applied across proteins.

### TMEM106B peptide level analysis (ROSMAP TMT proteomics)

TMEM106B peptide abundance in the two cohorts of brain ROSMAP TMT proteomics was tested for associations with common variation at *TMEM106B* (rs3173615), common variation at GRN (rs5848), and the interactive or additive effect between the two loci. An overlap check at the individual ID level identified five participants present in both datasets; these were removed from cohort 2, as cohort 2 was used for confirmatory analysis. The analysis then integrated genotype data, clinical covariates, biospecimen, sample-annotation resources, and TMEM106B-focused peptide quantification data derived from each dataset.

Genotypes for rs3173615 and rs5848 were linked to proteomic samples through the available sample-mapping framework and converted to additive dosage variables. rs3173615 was encoded as the number of G alleles (CC, CG, GG), and rs5848 was encoded as the number of T alleles (CC, CT, TT). Ambiguous genotype assignments at the individual level were excluded before analysis. TMEM106B peptide measurements were quality controlled by requiring Qvality q-value < 0.01, at least two peptide-spectrum matches, and removal of duplicate sequence-position entries. Across all analyses, peptides were divided into N-terminal and C-terminal groups on the basis of peptide start position relative to position 120, with C-terminal peptides considered as the proxy for the TMEM106B cleaved C-terminal fragment.

For each cohort, peptide-by-sample matrices were constructed after sample mapping. Non-positive intensities were treated as missing, values were log2-transformed, and sample-wise median centering was used for normalization. Peptides detected in at least 20% of mapped samples were retained for the primary analyses. TMEM106B peptides had extremely low percentages of missing values for C-terminal mapping peptides: SAYVSYDVQK had zero missing values and YQYVDCGR had only 15% missing values, and moderate percentages of missing values for N-terminal mapping peptides: DSVTCPTCQGTGR (23% of missing values), EDAYDGVTSENMR (43% of missing values), GQENQLVALIPYSDQR (35% of missing values), and NGLVNSEVHNEDGR (51% of missing values). Batch-wise MinProb imputation was not used during matrix construction, although limited row-wise low-intensity imputation was applied only at model fitting when sparse residual missing values remained. In practice, both imputation and non-imputation yielded similar findings.

Four prespecified models were fit in each cohort: a *TMEM106B* main-effect model, a *GRN* main-effect model, a *TMEM106B***×***GRN* interaction model, and a joint *TMEM106B* and *GRN* additive model without an interaction term. These analyses used linear modeling with limma empirical-Bayes moderation, with DEqMS-based spectrum-count-aware variance moderation applied when peptide-spectrum-match information was available. Models adjusted for major technical and biological covariates, including TMT batch where applicable, sex, age at death, and postmortem interval. Raw peptide-level *p* values were taken from the coefficient of interest, and false-discovery rates were controlled with the Benjamini-Hochberg procedure.

### Quantitation and statistical analysis

All quantification and statistical analyses were performed in R and Python. Analysis details can be found in figure legends, the methods, and the main text. All plots were prepared using tidyplots, ggplot2, ggpubr, and ggrepel in R.

